# Effects of larval and adult crowding on fitness components in *Drosophila* populations adapted to larval crowding experienced under different combinations of food amount and egg number

**DOI:** 10.1101/2022.05.08.491081

**Authors:** Neha Pandey, Rishabh Malhotra, Amitabh Joshi

**Affiliations:** Evolutionary Biology Laboratory, Evolutionary and Organismal Biology Unit, Jawaharlal Nehru Centre for Advanced Scientific Research, Jakkur, Bengaluru 560 064, India

**Keywords:** density-dependent selection, density-dependent feedback loops, population stability, life-history evolution, experimental evolution, *Drosophila melanogaster*

## Abstract

Since the realization in the 1970s that simple discrete-time population growth models can show complex unstable dynamics of population size, many explanations were proposed for the evolution of enhanced population stability. The most plausible of these was density-dependent selection, suggested to favour greater stability due to *r-K* trade-offs. However, the first experiment aimed at testing this prediction revealed that *Drosophila melanogaster* populations adapted to larval crowding did not evolve greater constancy stability than their ancestral controls. A subsequent study showed that *D. ananassae* populations adapted to larval crowding had evolved greater constancy and persistence than ancestral controls. These *D. ananassae* populations had experienced chronic larval crowding in conditions of very low amounts of food, whereas the earlier studied *D. melanogaster* populations had experienced chronic larval crowding at fairly high food amounts. Further theoretical work also suggested that populations adapting to crowding could evolve greater stability even in the absence of *r*-*K* trade-offs. Most recently, studies in our laboratory showed that two sets of crowding adapted *D. melanogaster* populations, derived from a common ancestral lineage, which differed in the food amounts at which they experienced larval crowding, evolved different patterns of constancy and persistence stability. These two sets of populations also differed in the traits, e.g. larval feeding rate, that evolved as they became more competitive. Here, we examine the response of key fitness components to larval and adult densities in these two sets of populations, to see whether differences in their stability attributes can be explained by variation in how their life-histories respond to crowding at different life stages. Of all traits examined, only pre-adult survivorship responded differently to larval density across the two sets of populations. The populations that adapted to larval crowding at low food amounts showed reduced sensitivity of pre-adult survivorship to larval density, compared to those that adapted to larval crowding at high food amounts. We discuss our results in the context of different ways in which density-dependent selection may facilitate the evolution of greater constancy or persistence, depending on the ecological details of how crowding was experienced.

## 1. Introduction

The recent focus on eco-evolutionary dynamics (e.g. Romero-Mujalli *et al* 2019; but see also Hendry 2019) notwithstanding, population ecology and evolutionary biology developed rather independently of one another during the first six to seven decades of the twentieth century (Kingsland 1995). Despite the early attempts of Elton (1927) to bridge the nascent fields of evolutionary biology and population ecology, pointing out that population cycles could have an effect on evolutionary dynamics because selection at high or low densities, respectively, might favour different sorts of traits, these two strands of density-dependent selection of varying traits and its possible effects on population dynamics and stability, began to come together only in the 1980s (reviewed in Mueller and Joshi 2000). On the one hand was the development of the formal theory of density-dependent selection (MacArthur 1962; MacArthur and Wilson 1967; Anderson 1971; Charlesworth 1971; Roughgarden 1971; Asmussen 1983), focussing attention on *r-K* trade-offs and their role in mediating adaptation to chronic crowding (Luckinbill 1978; Mueller and Ayala 1981; Mueller *et al* 1991; Vasi *et al* 1994); on the other, was the realization that simple population growth models could show varied and unstable dynamics of population size as long as intrinsic growth rates were high, and there was a time-lag in the density-dependent feedback (May 1974; May and Oster 1976), leading to investigations of proximal and ultimate causes of population stability (discussed in Jaggi and Joshi 2001; Mueller 2009; Dey and Joshi 2013, 2018).

Among the various proposed mechanisms for the evolution of population stability, the most plausible was that density-dependent selection could facilitate the evolution of stability, via an *r-K* trade-off, through promoting an evolved increase in *K* (reviewed by Mueller and Joshi 2000; Dey *et al* 2012). However, the first experiment aimed at testing this prediction did not support the notion that density-dependent selection would promote the evolution of greater stability in populations adapted to chronic crowding (Mueller *et al* 2000). Populations of *D. melanogaster* that had evolved traits indicating greater competitive ability than ancestral controls, as a result of having been subjected to high larval density for many generations, nevertheless showed no evidence of greater constancy stability (*sensu* Grimm and Wissel 1997) than controls, when reared for 68 generations in a food regime known to induce large and somewhat regular fluctuations in population size (first 45 generations reported by Mueller *et al* 2000; full study reported in Mueller and Joshi 2000). In this study, which involved very large cage populations (population size in the thousands), there were no extinctions and, hence, persistence could not be assessed. The lack of evolution of constancy was, however, not due to a general lack of evolutionary change, as other traits related to fitness under crowding did evolve in the populations studied (Mueller and Joshi 2000; Mueller *et al* 2000; Joshi *et al* 2003).

A subsequent study, using populations of *D. ananassae* adapted to larval crowding experienced under different ecological conditions than the populations used by Mueller *et al* (2000), on the other hand, showed clear evidence for density-dependent selection resulting in the evolution of enhanced stability (Dey *et al* 2012). The crowding adapted populations of *D. ananassae* exhibited significantly greater constancy and persistence than their ancestral control populations, but whether or not this was due to an underlying *r*-*K* trade-off was unclear (Dey *et al* 2012). While the crowding adapted populations had evolved both significantly higher average population size and estimated *K* than the controls, their estimated *r*, although substantially lower, was not significantly different from controls (Dey *et al* 2012). The sensitivity of realized per capita population growth rate to density was also significantly lower than controls in the crowding adapted populations (Dey *et al* 2012). The principal difference between the high larval crowding experienced by selected populations in the studies of Mueller *et al* (2000) and Dey *et al* (2012) was in the combination of food level and egg number in the culture vials. The populations of Mueller *et al* (2000) experienced larval rearing densities of ~1000-1500 eggs in 6-7 mL of banana-molasses food in 6-dram vials, whereas those of Dey *et al* (2012) were reared at larval densities of ~600 eggs in 1.5 mL cornmeal food in 8-dram vials. The larval crowding adapted populations of *D. ananassae* also evolved greater competitive ability via a different set of traits than the *D. melanogaster* populations of Mueller *et al* (2000): notably, the *D. ananassae* populations evolved greater time efficiency of larval food conversion to biomass, rather than an increased larval feeding rate (Nagarajan *et al* 2016). These differences led Dey *et al* (2012) to speculate that perhaps the precise suite of traits that evolves in response to chronic larval crowding may affect whether or not population stability evolves as a correlated response to density-dependent selection. However, any explanation for the differences between the results of Mueller *et al* (2000) and Dey *et al* (2012) could not be unequivocal, since the two studies also differed in species used, in addition to the food level and egg number combination at which they experienced larval crowding.

Subsequent selection experiments using a set of *D. melanogaster* populations (MCU populations: see Materials and Methods), derived from the same ancestors as those used by Mueller *et al* (2000), and adapted to larval crowding at an egg number and food level combination similar to that of the *D. ananassae* populations of Dey *et al* (2012), indicated that the specific suite of traits that evolved in response to chronic crowding was largely determined by egg number and food level combination, not by species identity (Sarangi *et al* 2016). Consequently, another set of *D. melanogaster* populations (LCU populations: see Materials and Methods), derived from the same ancestral controls as the MCU populations, and adapted to larval crowding at an egg number and food level combination similar to that of the *D. melanogaster* populations of Mueller *et al* (2000), was found to have evolved to adapt to chronic larval crowding via traits such as increased larval feeding rate, similar to those evolved by the populations used by Mueller *et al* (2000) (Sarangi 2018). As a result of these various studies, it became clear that the LCU and MCU populations, which shared ancestral controls, evolved to adapt to chronic larval crowding via different suites of traits as a result of the different combinations of egg number and food level at which they experienced crowding (Sarangi 2018; Venkitachalam *et al* 2022). These studies, thus, paved the way for testing the speculative predictions made by Dey *et al* (2012) about whether the specific traits that evolve in response to chronic larval crowding could affect whether or not population stability evolves as a correlated response to density-dependent selection.

To this end, the dynamics and stability of the LCU and MCU populations, relative to their ancestral controls, were assessed in two separate multi-generation population dynamic experiments that revealed that the MCU populations, like the *D. ananassae* populations of Dey *et al* (2012) had evolved greater constancy and persistence than controls, whereas the LCU populations, like those of Mueller *et al* (2000) did not show greater constancy than controls, but did exhibit higher persistence (N Pandey and A Joshi, *unpubl. mss*.). The MCU populations also exhibited greater average population size and estimated *K*, and reduced sensitivity of realized per capita population growth rate to density, as compared to controls (N Pandey and A Joshi, *unpubl. ms*.). The LCU populations, on the other hand, did not differ from controls in average population size and estimated *K*, but did show significantly lower density-sensitivity than controls of realized per capita population growth rate (N Pandey and A Joshi, *unpubl. ms*.). It was this observed difference in the dynamics and stability characteristics of the MCU and LCU populations that motivated the present study which examined whether the MCU and LCU populations differed in major life-history traits and their sensitivity to density.

## 2. Materials and methods

In this study, since we were interested in examining the density-responses of life-history traits in the context of their role in mediating population dynamics, we used food amounts and egg numbers that approximate the conditions in typical population dynamics experiments in our laboratory. Consequently, the larval density conditions in the present assays are different from previous assays of life-history traits in these populations under low and high larval densities (e.g. Sarangi *et al* 2016; Sarangi 2018; Venkitachalam *et al* 2022).

### 2.1. Experimental populations

We used eight large outbred laboratory populations of *D. melanogaster* for this study, belonging to two selection regimes, consisting of four replicate populations each, subjected to larval crowding at different combinations of egg number and food amount. Complete details of the derivation and maintenance of these populations can be found in Sarangi (2018); we reiterate the essential details here.

#### 2.1.1. MCU 1-4

Each MCU population was derived from an independent ancestral population, and they were maintained at a high density of around 600 eggs in 1.5 mL of cornmeal food, in cylindrical Borosilicate glass vials of 2.2-2.4 cm inner diameter and 9.5 cm height. The MCUs had been maintained on a 21-day discrete-generation cycle, under constant light (LL), at 25° ± 1°C temperature and around 80% humidity for about 185 generations at the time of this study. To maintain a breeding population size of around 1800 adults per vial, 12 vials per population were set up for each generation. Since larval crowding prolongs development, eclosing adults were collected into cages every day till day 18 from egg-lay, once eclosions began. Eclosing adults from all vials of a population were transferred to a Plexiglas cage (25 × 20 × 15 cm^3^) containing a food plate (Petridish with cornmeal food), and a wet cotton ball to help maintain humidity. The old food plate was replaced with a fresh food plate every alternate day till day 18 from egg-lay, and the cotton ball was changed at every alternate food plate change. On the 18^th^ day, adults were given live acetic-acid-yeast supplement, in addition to the regular food, till day 20, and on the 20^th^ day these adults were allowed to lay eggs for around 18 hours on vertically cut sterile cornmeal food. On the 21^st^ day, eggs laid by these flies were collected into vials to start the next generation.

#### 2.1.2. LCU 1-4

Each LCU population was derived from an independent ancestral population, the same set of four ancestors as the MCUs, and they had been maintained at a high larval density of around 1200 eggs in 6 mL of cornmeal food per vial for about 64 generations at the time of this study. During the pre-adult stage, LCUs were maintained in slightly shorter and narrower vials (9 cm height × 2.0-2.2 cm inner diameter) than MCUs. The egg number, food amount, and vial dimensions of the LCUs were chosen to approximate the maintenance regime of the CUs at the larval stage (first described in Mueller *et al* 1993) used in earlier studies. Except for the difference in egg number and food amount, and vial dimensions, the maintenance protocol for LCUs was the same as for MCUs.

### 2.2. Trait assays

Prior to assays, we subjected all populations to one generation of common rearing at low larval density (roughly 70 eggs in 6 mL cornmeal food per vial), corresponding to that used for the ancestral control populations for both the MCU and LCU populations, to avoid non-genetic parental effects contributing to differences among selection regimes. After a generation of common rearing, we allowed females to lay eggs for 12 hours on a sterile double agar plate, and these eggs were used to initiate the various assays. Due to logistical constraints, only two replicate populations per selection regime were assayed at a time: MCU 1-2 (184 generations of selection) and LCU 1-2 (63 generations of selection) were assayed together, and MCU 3-4 (185 generations of selection) and LCU 3-4 (64 generations of selection) were assayed together.

#### 2.2.1. Pre-adult survivorship

We examined the pre-adult survivorship for all the selected and control populations in the following egg number × food amount combinations: (i) 75 eggs in 1 mL and 1.5 mL cornmeal food (10 vials each); (ii) 150 eggs in 1 mL and 1.5 mL cornmeal food (5 vials each); (iii) 300 eggs in 1 mL and 1.5 mL cornmeal food (5 vials each). We counted all the eclosing adults to calculate pre-adult survivorship in each vial.

#### 2.2.2. Female fecundity

We collected flies eclosing between days 9-11 after egg-lay in the vials kept at different population × egg number × food amount combinations for the pre-adult survivorship assay. These flies were kept in groups of five males and five females per vial, containing 4 mL cornmeal food), with 5 such vials set up for each selection regime × egg number × food amount combination on days 9, 10 and 11 from egg-lay (following Vaidya 2013). These collected flies were shifted to fresh food vials every alternate day till day 17 from egg-lay. Prior to assaying fecundity on day 21 post egg-lay, the flies were given live acetic-acid-yeast paste supplement in the food vials from day 18 post egg-lay. On day 20 from egg-lay, we mixed all flies from all the 15 vials for each selection regime × egg number × food amount combination, and from this pool of flies that varied in adult age from 10-12 days old, we haphazardly chose 15 males and 15 females for assaying fecundity. Individual male-female pairs were placed into a sterile food vial containing a thin layer of food for egg laying over the next 16 hours, after which we removed the flies and counted the number of eggs under a stereo-zoom microscope. Thus a total of 720 females (8 populations × 6 treatment combinations × 15 replicates) were assayed for fecundity on day 21 post egg-lay.

#### 2.2.3. Dry body-weight at eclosion

We thoroughly mixed the flies eclosing in vials kept at different population × egg number × food amount combinations for the pre-adult survivorship assay, and then haphazardly chose 25 males and 25 females per population × treatment combination for weighing. We refrigerated these flies at 5°C, and later dried them for 36 hours at 70°C in a hot air oven before weighing them to the nearest 10^-5^ g on a Sartorius CP225D microbalance, in single-sex groups of five flies each. Each batch of five flies was weighed thrice, and the mean of those readings taken as the dry bodyweight of that batch.

#### 2.2.4. Adult survivorship till egg-lay

We examined the effect of adult crowding on adult survivorship from eclosion till day 21 post egg-lay, the last day of effective adult life in three-week discrete-generation cultures, by subjecting the adults to two densities. We reared larvae at a low larval density of ~70 eggs in 6 mL cornmeal food per vial. On day 11 from egg-lay, we shifted eclosing adults into one of two adult density treatments: low (50 adults per vial) and high (150 adults per vial), with a 1:1 sex ratio in each vial. We had exactly 5 mL of cornmeal food in all the vials and used cotton plugs of similar size to plug the vials so as to offer equal space for flies in all vials. Each adult density treatment had 7 replicate vials per population. Flies were moved to fresh food vials every day to avoid confounding effects of larval activity. The number of flies dying in each vial per day, till day 21 post egg-lay was recorded.

### 2.3. Statistical analyses

Every replicate larval crowding adapted population shares ancestry with an MB population with the same replicate subscript i.e. replicate population *i* in the MCU, and LCU regimes is derived from replicate *i* of MB (*i* = 1..4). This permits the use of a completely randomized block design in our statistical analysis, with replicate populations bearing the same subscript treated as blocks. In the mixed model analyses of variance (ANOVAs), block was treated as a random factor crossed with the fixed factors selection regime, egg number, and food amount. An additional fixed factor, sex, was also added while analyzing dry body-weight data. For testing the effect of adult density on adult survivorship, block was treated as random factor, and selection regime, adult density, and sex were treated as fixed factors. We performed all analyses on Statistica Version 5.0 (Statsoft 1995), and used Tukey’s HSD at *α* = 0.05 for all post-hoc multiple comparisons.

We also analyzed the data on pre-adult survivorship, female fecundity at day 21 post-egg lay, and male and female dry body-weight at eclosion via linear regression against larval density (eggs per unit volume food) as the independent variable, utilizing the fact that the six combinations of egg number and food amount corresponded to six different larval densities. Linear regression for each replicate MCU and LCU population was carried out separately, and then the eight values obtained for slope, intercept and *R^2^*, respectively, were subjected to mixed model ANOVA with selection regime as a fixed factor and block as a random factor. All these analyses were also implemented in Statistica Version 5.0 (Statsoft 1995).

## 3. Results

### 3.1. Pre-adult survivorship

As expected, pre-adult survivorship was lower at higher egg numbers, in both 1 mL and 1.5 mL food, for both LCU and MCU populations (Figure 1 A,B). For both sets of populations, the drop in survivorship was greater between 150 and 300 eggs per vial, compared to from 75 to 150 eggs (Figure 1 A,B). There was slightly lower survivorship in 1 mL than in 1.5 mL (Figure 1 A,B), but there were no significant effects of food amount or any interaction involving food amount (Table 1). The only significant ANOVA effects were that of egg number and the selection × egg number interaction (Table 1), driven by significantly higher survivorship of MCU over LCU populations, by about 20%, at the highest egg number of 300 per vial (Tukey’s HSD, *P* < 0.05), in both the 1 mL and 1.5 mL food amount treatments (Figure 1 A,B).

**Figure 1:**
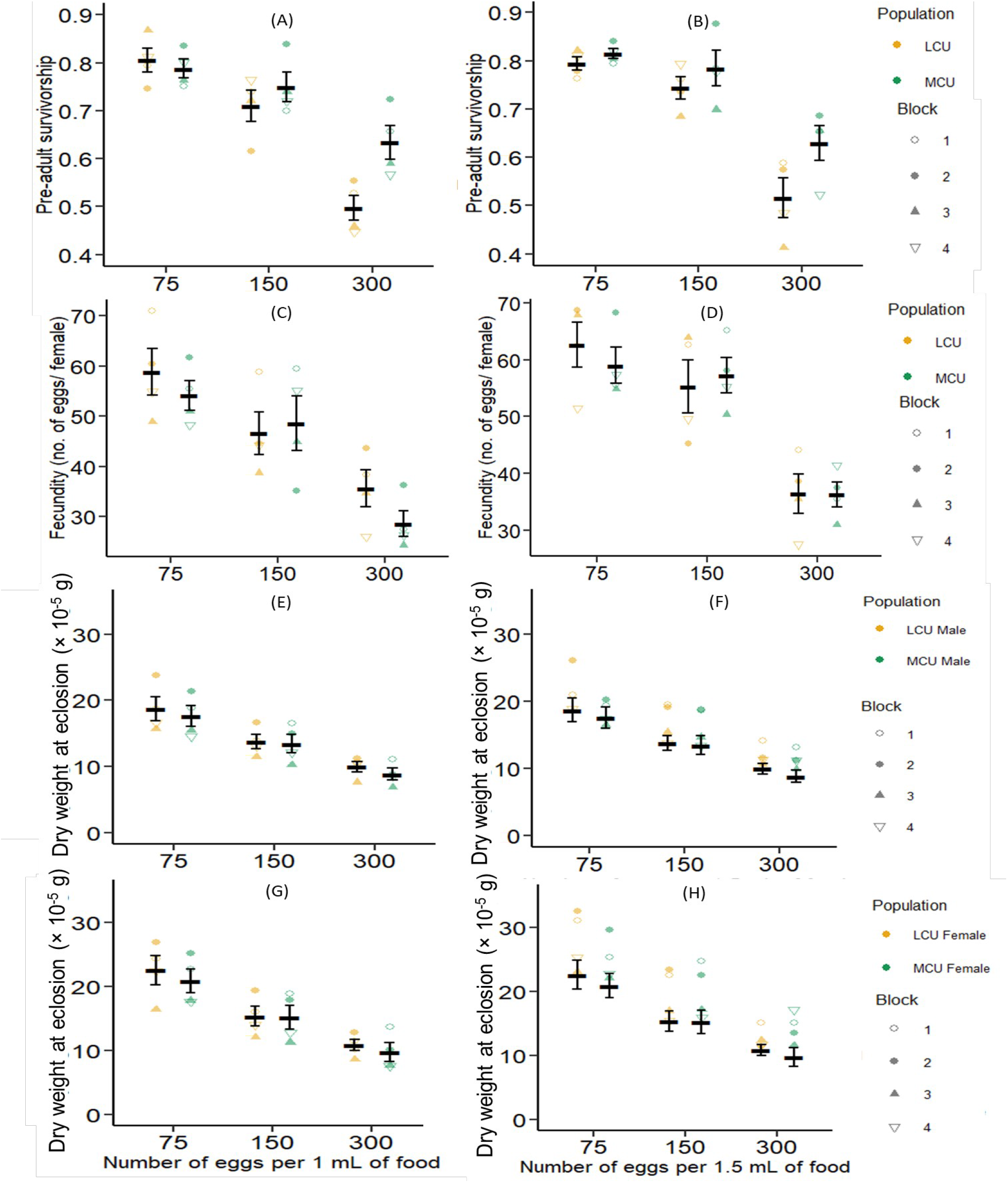
Mean pre-adult survivorship, averaged over the four replicate populations each, of the crowding-adapted MCU and LCU populations at different egg numbers and larval food volume in 1 mL (A) and in 1.5 mL (B). Mean fecundity, averaged over the four replicate populations each, of the crowding-adapted MCU and LCU populations at different egg numbers and larval food volume in 1 mL (C) and in 1.5 mL (D). Mean dry body-weight at eclosion for males, averaged over the four replicate populations each, of the crowding-adapted MCU and LCU populations at different egg numbers and larval food volume in 1 mL (E) and in 1.5 mL (F). Mean dry body-weight at eclosion for females, averaged over the four replicate populations each, of the crowding-adapted MCU and LCU populations at different egg numbers and larval food volume in 1.5 mL (G) and in 1 mL (H). Error bars around the means are standard errors based on the means of four replicate populations within each selection regime.

**Table 1.** Results from ANOVA done on pre-adult survivorship. Since we were primarily interested in fixed main effects and interactions, block effects and interactions have been omitted for brevity.

| Effect | *df* | MS | *F* | *P* |
| --- | --- | --- | --- | --- |
| Selection | 1 | 0.224897 | 4.438 | 0.125 |
| Egg number | 2 | 1.446999 | 34.781 | <0.001** |
| Food amount | 1 | 0.018818 | 2.852 | 0.190 |
| Selection × Egg number | 2 | 0.102034 | 5.373 | 0.046* |
| Selection × Food amount | 1 | 0.000556 | 0.047 | 0.842 |
| Egg number × Food amount | 2 | 0.005316 | 1.068 | 0.401 |
| Selection × Egg number × Food amount | 2 | 0.006928 | 1.640 | 0.270 |

\*: $P < 0.05$ ; \*\*: $P < 0.01$

### 3.2 Female fecundity at day 21 post egg-lay

As expected, female fecundity on day 21 post egg-lay was lower at higher egg numbers, and in 1 mL compared to 1.5 mL food, for both LCU and MCU populations (Figure 1 C,D). ANOVA revealed that only the main effects of egg number and food amount were significant (Table 2). For both sets of populations, the drop in fecundity was greater between 150 and 300 eggs per vial, compared to from 75 to 150 eggs (Figure 1 C,D). Fecundity of LCU females (mean: 35.6 eggs) was visibly greater than that of MCU females (mean 28.6 eggs) in the combination of 300 eggs in 1 mL food (Figure 1 C), but it was not possible to make anything of it in the absence of any significant interaction of selection × egg number × food amount.

**Table 2.** Results from ANOVA done on female fecundity on day 21 post egg-lay. Since we were primarily interested in fixed main effects and interactions, block effects and interactions have been omitted for brevity.

| Effect | *df* | MS | *F* | *P* |
| --- | --- | --- | --- | --- |
| Selection | 1 | 657.4 | 0.528 | 0.520 |
| Egg number | 2 | 38135.0 | 27.681 | <0.001** |
| Food amount | 1 | 6008.9 | 12.802 | 0.037* |
| Selection × Egg number | 2 | 684.5 | 3.611 | 0.093 |
| Selection × Food amount | 1 | 309.4 | 0.388 | 0.578 |
| Egg number × Food amount | 2 | 377.8 | 0.733 | 0.519 |
| Selection × Egg number × Food amount | 2 | 215.4 | 0.430 | 0.669 |

\*: $P < 0.05$ ; \*\*: $P < 0.01$

### 3.3 Dry body-weight at eclosion

Male (Figure 1 E,F) and female (Figure 1 G,H) dry body-weights at eclosion showed the expected prominent effects of sex (females heavier), egg number and food amount (weight decrease with increase in egg number and reduction in food), but there was no clear indication of any difference in how the two sets of populations responded to increasing egg number or decreasing food amount (Figure 1 E,F,G,H). Correspondingly, in the ANOVA, the main effects of sex, egg number and food amount were significant (Table 3). Male dry weights (Figure 1 E,F) were less severely affected by increasing larval density than female dry weights (Figure 1 G,H), driving significant egg number × sex and food amount × sex interactions (Table 3). There was no main effect of selection regime, and only one interaction involving selection regime (selection × egg number × food amount) was significant (Table 3). However, this interaction was due to small haphazard differences in the response of how dry body-weights changed in LCU and MCU populations across egg number and food amount combinations, and these changes did not suggest any clear pattern of differences between selection regimes in the sensitivity of dry bodyweight to larval density (Figure 1 E,F,G,H).

**Table 3.**
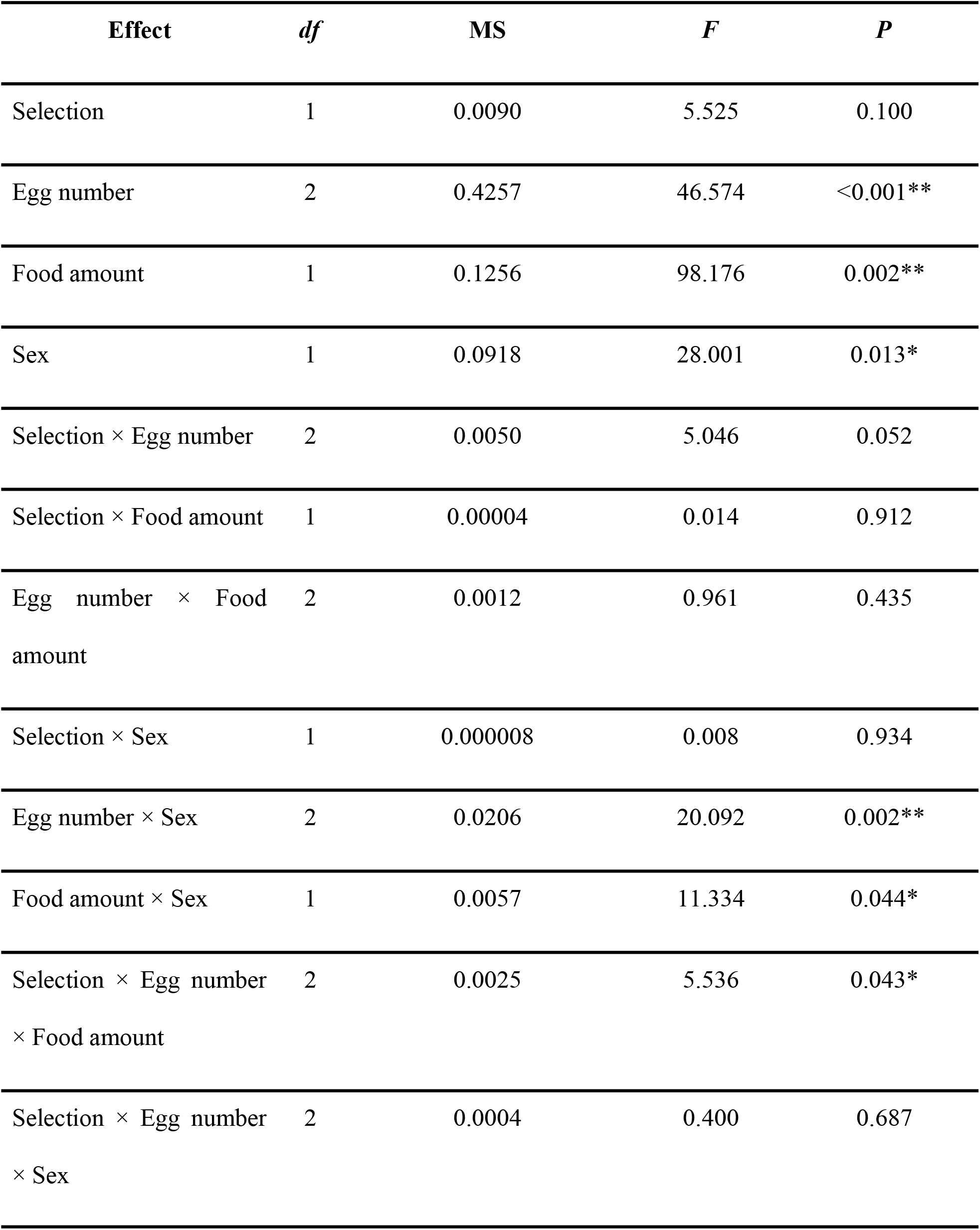
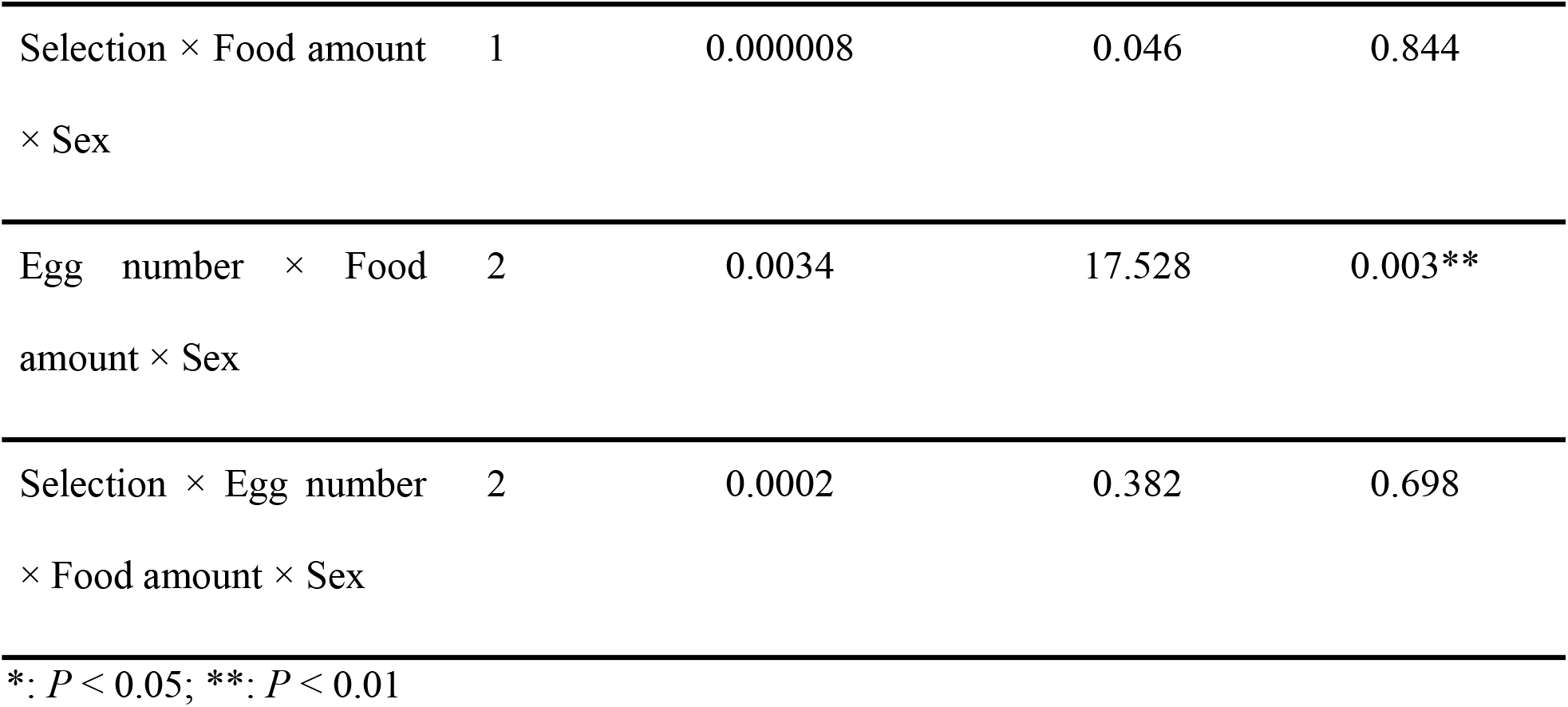
Results from ANOVA done on dry body-weight per fly (in mg) at eclosion. Since we were primarily interested in fixed main effects and interactions, block effects and interactions have been omitted for brevity.

### 3.4 Adult survivorship till egg-lay

As expected, adult survivorship from eclosion to day 21 post-egg lay declined sharply as density increased from 50 flies per vial (mean survivorship: 0.94) to 150 flies per vial (mean survivorship: 0.48) (Figure 2 A,B). Female (Figure 2 B) and male (Figure 2 A) survivorship was somewhat similar at 50 flies per vial (mean survivorship females: 0.90, males: 0.97), whereas at 150 flies per vial, females had markedly lower mean survivorship (0.29) than males (0.67). These results were reflected in significant ANOVA effects of sex, adult density, and the sex × adult density interaction (Table 4). There was no evidence for any difference between selection regimes in the response of adult survivorship to adult density (Figure 2 A,B), and neither the main effect of selection regime, nor any interaction involving selection regime, was significant (Table 4).

**Figure 2:**
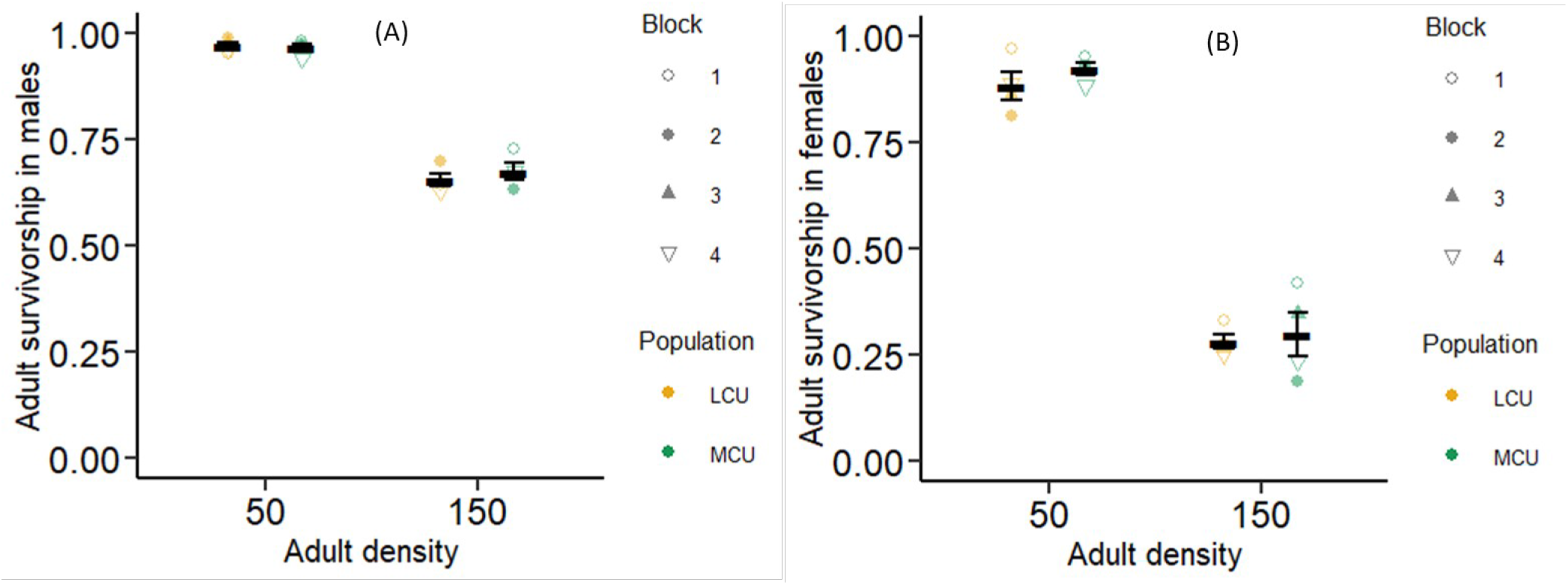
Mean adult survivorship, averaged over the four replicate populations each, of males (A) and females (B) from the crowding-adapted MCU and LCU populations at low (50 adults) and high (150) adult density. Error bars around the means are standard errors based on the means of four replicate populations within each selection regime.

**Table 4:** Results from ANOVA done on adult survivorship from eclosion to day 21 post egg-lay. Since we were primarily interested in fixed main effects and interactions, block effects and interactions have been omitted for brevity.

| Effect | *df* | MS | *F* | *P* |
| --- | --- | --- | --- | --- |
| Selection | 1 | 0.0172 | 1.634 | 0.291 |
| Adult density | 1 | 11.8435 | 2086.225 | <0.0001** |
| Sex | 1 | 2.7428 | 84.928 | 0.003** |
| Selection × Adult density | 1 | 0.00001 | 0.000 | 0.988 |
| Selection × Sex | 1 | 0.0055 | 0.665 | 0.475 |
| Adult density × Sex | 1 | 1.3558 | 405.599 | <0.001** |
| Selection × Adult density × Sex | 1 | 0.0075 | 1.061 | 0.379 |

\*: $P < 0.05$ ; \*\*: $P < 0.01$

### 3.5 Regression analyses

The results from the regression analyses were concordant with those described above in Sections 3.1 – 3.3 (Figure 3). There were no significant differences seen between MCU and LCU populations, on an average, for either intercept or goodness of fit (*R*^2^) for pre-adult survivorship, female fecundity on day 21 post-egg lay, or male/female dry body-weight at eclosion (ANOVA results not shown). Only one of the ANOVAs on slope showed a significant difference between selection regimes (*F*_1,3_ =15.91, *P* = 0.028); the MCU populations (Figure 3 A) had a significantly less negative slope than the LCU populations (Figure 3 B) for pre-adult survivorship. Thus, MCU populations had a higher pre-adult survivorship than LCU populations over a wide range of larval densities. For female fecundity (Figure 3 C,D), and dry body-weight of males (Figure 3 E,F) and females (Figure 3 G,H), the slopes did not significantly differ between selection regimes (ANOVA results not shown).

**Figure 3:**
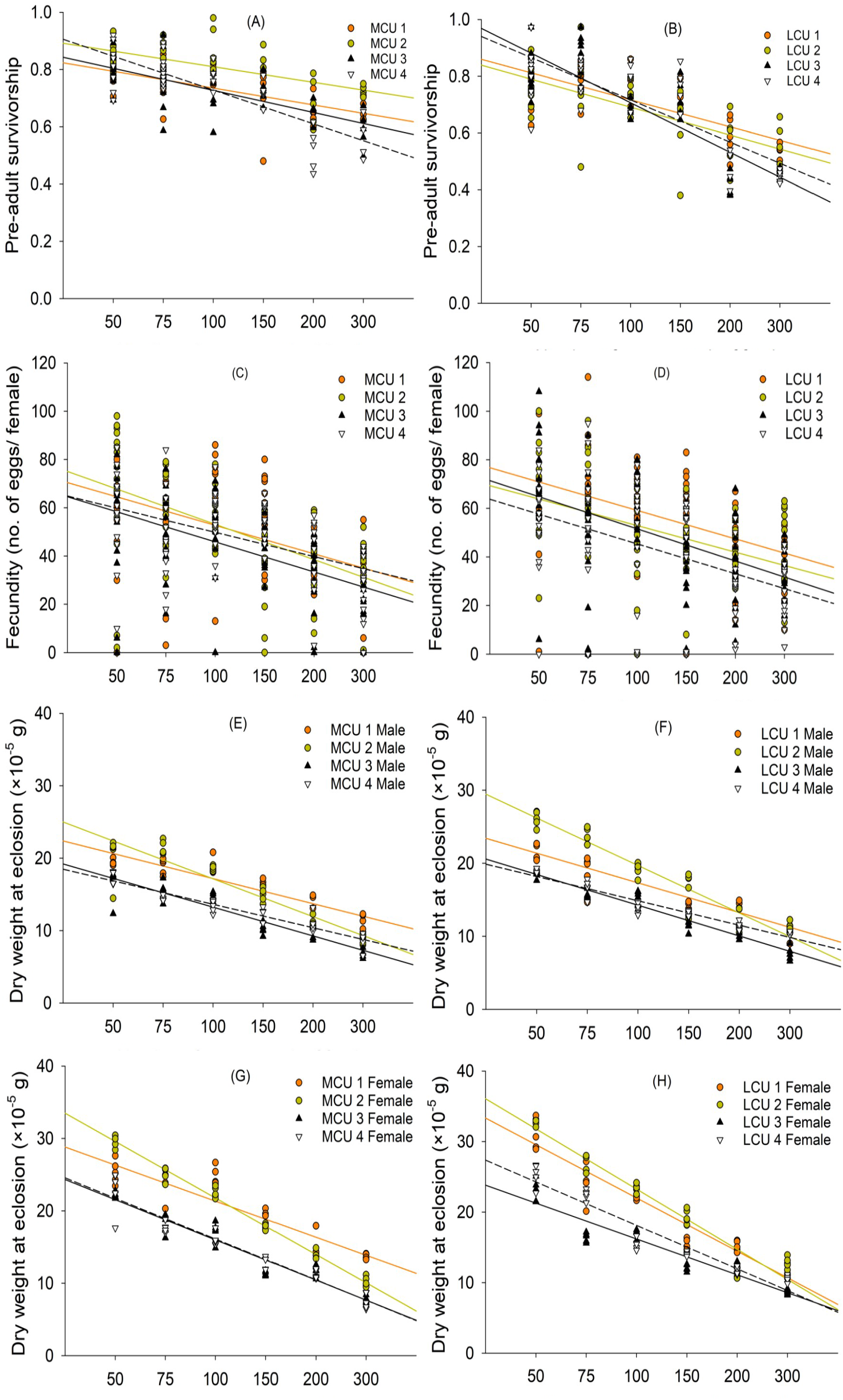
Pre-adult survivorship regressed on egg density for replicates of the crowding-adapted MCU (A) and LCU (B) populations. Fecundity regressed on egg density for replicates of the crowding-adapted MCU (C) and LCU (D) populations. Dry body-weight for males regressed on egg density for replicates of crowding-adapted MCU (E) and LCU (F) populations. Dry body-weight for females regressed on egg density for replicates of crowding-adapted MCU (G) and LCU (H) populations.

## 4. Discussion

We undertook this study to examine whether differences between the MCU and LCU populations in the sensitivity of key life-history traits to larval or adult density might explain the fact that greater constancy evolved in the MCU, but not the LCU populations (N Pandey and A Joshi, *unpubl. mss*.), as a consequence of density-dependent selection due to chronic larval crowding. In *Drosophila* cultures, there are four major density-dependent negative-feedback loops that affect population growth: larval density-dependent larval mortality, larval density-dependent adult size and, therefore, female fecundity, adult density-dependent adult mortality, and adult density-dependent female fecundity (Figure 4; Mueller 1988; Joshi *et al* 1998). In LH food regimes, the sensitivity of female fecundity to adult density is considerably weakened due to the provision of supplementary yeast past, resulting in cyclic dynamics (Mueller and Huynh 1994), and detailed individual based simulations indicate that this feedback loop has relatively little effect in further modulating the dynamics of cultures in an LH food regime (Tung *et al* 2019). Consequently, we examined only the sensitivity to density of pre-adult survivorship, female fecundity at day 21 post-egg lay, male and female dry weight at eclosion, and adult mortality from eclosion to day 21 post-egg lay in the MCU and LCU populations.

**Figure 4:**
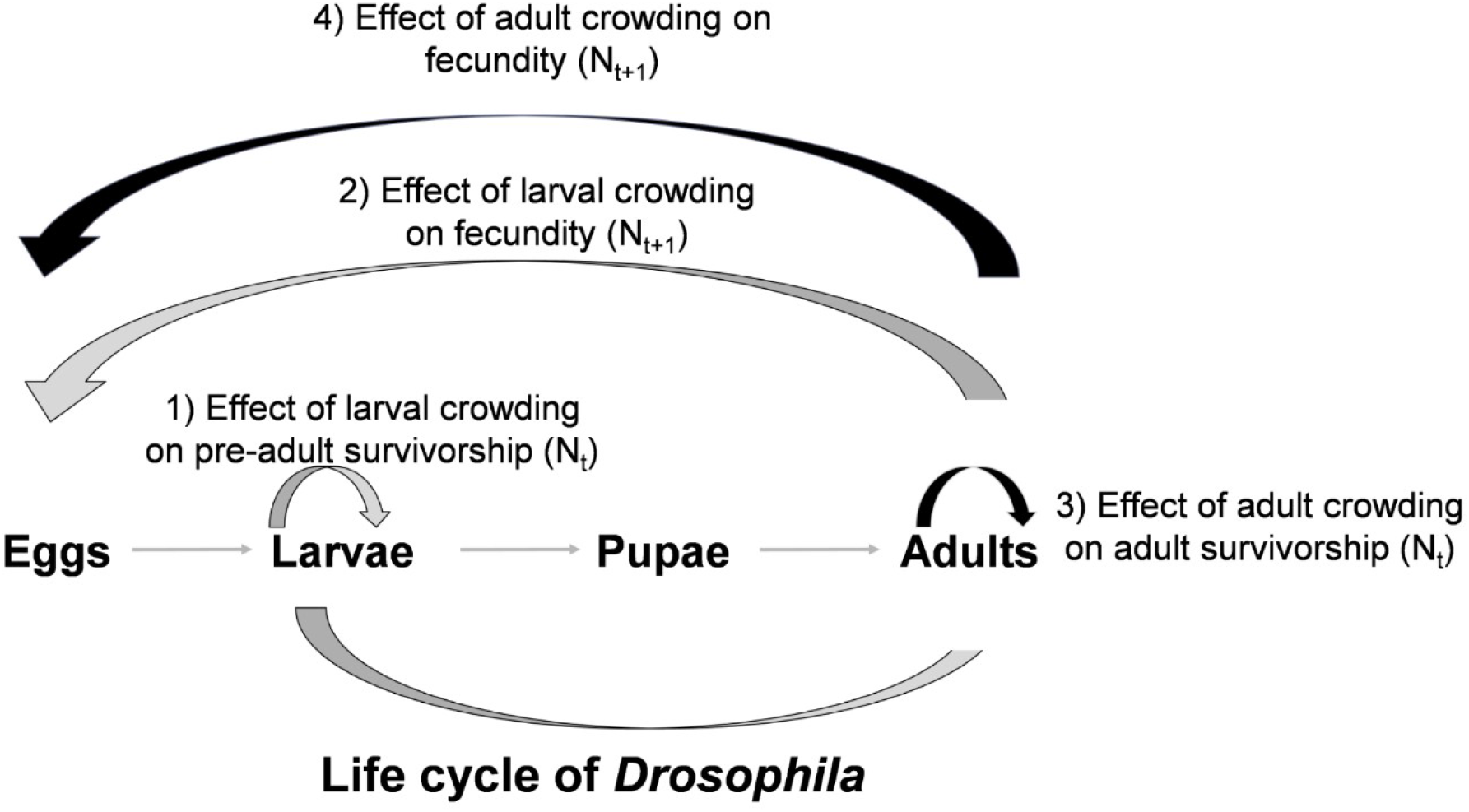
Life cycle of *Drosophila melanogaster*, and the four density-dependent feedback loops operating at different life-stages: (1) Effect of larval crowding on pre-adult survivorship, (2) Effect of larval crowding on female fecundity, (3) Effect of adult crowding on adult survivorship, and (4) Effect of adult crowding on female fecundity. The grey arrows show the effects of larval density and the black arrows show the effect of adult density.

To briefly recapitulate, earlier studies (N Pandey and A Joshi, *unpubl. mss*.) showed that both the MCU and the LCU populations showed significantly greater persistence, and significantly lower sensitivity of realized per capita growth rate to density, than their common controls. In addition, the MCU populations also showed significantly greater constancy, equilibrium population size, and average population size than controls. The LCU populations had very similar constancy to controls, and their average population size and equilibrium population size were higher than controls, but not significantly so. In terms of the magnitude of difference from controls, the MCU populations differed from controls, on an average, to a degree greater than the LCU populations did, for both persistence and the sensitivity of realized per capita growth rate to density (N Pandey and A Joshi, *unpubl. mss*.).

In the present study, the only life-history trait that differed significantly in its sensitivity to density between the MCU and LCU populations was pre-adult survivorship (Figures 1 A,B and 3 A,B). This particular density-dependent feedback loop is a large contributor to stability, with decreased sensitivity of pre-adult survivorship to larval density tending to stabilize the dynamics with regard to constancy (Mueller 1988; Mueller and Joshi 2000). In discrete-generation populations, greater pre-adult survivorship at a wide range of larval densities can potentially enhance both constancy and persistence by raising the population size at troughs in populations undergoing large fluctuations in population size due very strong time-lagged density-dependent feedback. It is therefore likely that the significantly and markedly lower sensitivity of pre-adult survivorship to larval density in the MCU populations is the major contributor to the evolution of enhanced constancy in the MCU but not the LCU populations. We speculate that the evolution of greater persistence than controls in both the MCU and LCU populations is again driven by a lower sensitivity of pre-adult survivorship to larval density, as reflected in the earlier observed ability of both MCU and LCU populations to sustain a higher realized per capita growth rate across medium to high adult densities (N Pandey and A Joshi, *unpubl. mss*.).

We know that adaptation to larval crowding at a much lower amount of food as compared to the LCU populations has led to the MCU populations evolving to attain their minimum critical size faster than controls; the LCU populations, on the other, evolved a higher feeding rate than controls (Sarangi *et al* 2016; Sarangi 2018). We speculate that during the population dynamics experiment, the faster attainment of minimum critical size may be driving increased pre-adult survivorship in the MCU populations, as larval cultures in such experiments are likely to quickly become toxic due to accumulation of metabolic waste in very low food amount (1/1.5 mL LH food in population dynamics experiments). On the other hand, the LCU populations, which have evolved a higher feeding rate, may be undergoing greater mortality due to metabolic waste buildup in the food; faster feeding is likely to be harmful under high concentrations of metabolic waste (Mueller and Barter 2015).

Overall, the present study, together with earlier population dynamics studies on these populations (N Pandey and A Joshi, *unpubl. mss*.), provide empirical support for the speculation by Dey *et al* (2012) that the differing suites of traits that evolve in response to chronic larval crowding experienced at different combinations of egg number and food amount are likely to differentially affect equilibrium population size and the sensitivity of realized per capita growth rate to density, thereby potentially affecting population stability. What ultimately links individual traits like feeding rate, minimum food requirement for development, or metabolic waste tolerance to demographic attributes like realized density-dependent per capita growth rates under competitive conditions is the sensitivity of key life-history traits to density. This study is the first attempt to connect trait evolution during density-dependent selection, experienced under differing ecological contexts, to the dynamics and stability of populations via the sensitivity of life-history traits to density. We suggest that these linkages need to be explored further, both empirically and theoretically, in a model-free framework, in order to achieve a better understanding of when and how density-dependent selection affects the evolution of population dynamics and stability attributes.

## Acknowledgments

We thank Sajith V. S., Ennya Anna Thomas and Srikant Venkitachalam for help in the assays, and Avani Mital, Manaswini Sarangi, N. Rajanna and Muniraju for assistance in the maintenance of populations. NP was supported by a doctoral fellowship from the Jawaharlal Nehru Centre for Advanced Scientific Research. RM’s stay in the lab was supported by Summer Research Fellowship from the Jawaharlal Nehru Centre for Advanced Scientific Research. This work was supported by a J.C. Bose National Fellowship (SERB, Government of India) to AJ and, in part, by AJ’s personal funds.

